# Shared TCRs in peripheral blood offer robust celiac disease classification independent of gluten Intake

**DOI:** 10.64898/2025.12.26.696039

**Authors:** Rebecca Elyanow, Rok Seon Choung, Eric V. Marietta, Ramit Bharanikumar, Wenyu Zhou, Haiyin Chen-Harris, Govind Bhagat, Peter H. Green, Bryan Howie, Harlan S Robins, Susan L. Neuhausen, Joseph A. Murray

## Abstract

**Background and Aims:** Celiac disease (CeD) is a chronic digestive autoimmune disorder affecting approximately 1% of the worldwide population. It is driven by T cells activated by specific HLA-DQ2 or HLA-DQ8 molecules leading to the destruction of intestinal villi. We aimed to characterize shared CeD-specific T cells in patients on and off a gluten-free diet (GFD) from a large cohort of cases and controls.

**Methods:** We performed bulk TCRβ immune sequencing of the peripheral blood of 1,604 biopsy-confirmed CeD patients (1,339 on GFD, 265 on normal diet) and 1,100 controls. We identified over 300 TCRβs enriched in CeD cases versus controls in an HLA-aware manner, controlling for CeD risk alleles.

**Results:** CeD-associated TCRβs were found to be more predictive of disease than previously characterized gliadin and glutenin-binding TCRs in a validation cohort. Furthermore, the clonal breadth of these TCRβs was associated with increased intestinal damage. Immune sequencing of the peripheral blood also uncovered repertoire-level differences between CeD patients and controls. CeD patients displayed significantly higher productive clonality compared to age-matched controls as well as expansion of TCRβs specific to cytomegalovirus (CMV) and Epstein-Barr Virus (EBV).

**Conclusions:** These findings underscore the value of unbiased immune repertoire sequencing to identify novel biomarkers for autoimmune disease and to discover new disease mechanisms which can improve both diagnosis and treatment of disease.

## Introduction

Celiac disease (CeD) is an inflammatory gastrointestinal disease caused by the ingestion of gluten [1,2]. In individuals with CeD, consumption of gluten results in enteropathy, and a gluten-free diet is the standard treatment [3]. It is one of the most common chronic digestive disorders, affecting approximately 1% of the population worldwide, and the prevalence of the disease continues to increase [4]. CeD occurs only in individuals with susceptible genes, the class II human leukocyte antigen (HLA) alleles, HLA-DQ2 and HLA-DQ8 [1,5]. HLA-DQ2.5 confers the greatest celiac disease risk, with DQ2.2 and DQ7.5 forming DQ2.5 in trans configuration, further contributing to susceptibility [6]. In CeD, both DQ2 and DQ8 present gluten-derived peptides to CD4+ T cells as antigens [7]. Numerous studies have demonstrated that several gluten-derived peptides can stimulate CD4+ T cells in patients with CeD, most notably the 33-mer alpha-gliadin peptide with 6 overlapping immunogenic epitopes [8, 9]. These anti-gluten CD4+ T cells and associated upregulation of IL-15 provide intraepithelial lymphocytes (CD8+) with a “license to kill” intestinal epithelial cells, resulting in tissue destruction in the small bowel [9,10,11,12].

Despite excellent evidence for the causal role of gluten-derived antigens, several puzzles remain in understanding the pathogenesis of CeD. For example, what triggers an increase in antibodies to the self-antigen tissue transglutaminase in the absence of a strong response by T cells to gluten-derived foreign antigens [13,14]? And why do many individuals with HLA risk alleles never develop CeD? Exogenous antigens other than gluten, such as viral infections, especially gastrointestinal viruses, have been hypothesized to play a role in triggering disease, but no causal antigens or T cell receptors (TCRs) have been discovered [15]. While the gluten-specific CD4+ T cell is central to the pathogenesis of CeD, it is highly likely that T-cell responses to other self or exogenous antigens also occur in CeD and are important for disease progression. To understand the causal mechanism behind CeD, it is crucial to identify additional TCRs associated with CeD beyond the known TCRs responding to gluten.

Next generation sequencing of adaptive immune repertoires and advanced analytics have provided researchers with an unbiased approach to determining underlying TCRs involved in disease-specific T cell responses [16,17]. Several studies have identified disease-specific TCRs by comparing the T cell repertoire of a given disease cohort with that of control cohorts [18,19]. Prior studies have performed immune sequencing of celiac patients, but they are either restricted to tissue-specific T cells or gliadin-stimulated T cells [20-25]. In this study, we analyzed the peripheral circulating TCR repertoire of CeD patients, from both treated (gluten-free diet) and untreated healthy controls. This study represents the largest immune sequencing effort of the peripheral blood of CeD patients as well as the largest of treated CeD patients.

The diversity of the T cell immune repertoire is critical in maintaining an effective immune response. Several studies have shown that a decrease in T cell diversity is common to several autoimmune diseases, such as rheumatoid arthritis, multiple sclerosis, and type 1 diabetes [26-29]. However, the diversity of the TCR repertoire in CeD had not been previously studied in an unbiased way. We postulated that CeD patients would display a less diverse TCR repertoire compared to controls. Moreover, we hypothesized that some specific, public, TCRs may react to antigens beyond known targets.

In this study, we identified 370 shared TCRs specific to CeD patients but not controls, including 318 novel CeD-associated TCRs. We demonstrated that these novel TCRs were more diagnostic of CeD than previously characterized gliadin-specific TCRs, particularly for subjects adhering to a gluten-free diet. We additionally investigated whether CeD induced large-scale changed to the peripheral T cell repertoire and found that CeD patients had significantly less diverse repertoires than controls and exhibited clonal expansion of T cells specific to common viral exposures.

## Materials and Methods

### Study Population

Our study population comprised cohorts from three distinct study populations across the US and Canada, including: 613 celiac disease (CeD) patients and 696 controls from the Beckman Research Institute of City of Hope (BRICOH), 728 CeD patients and 147 controls from the Mayo Clinic (Rochester, MN), and 263 CeD patients and 257 controls from the Seattle area collected by Adaptive Biotechnologies. All CeD patients were confirmed via duodenal biopsy; controls have not been biopsy confirmed negative.

Our validation cohort included 8032 previously sequenced healthy donors and donors with other infectious and autoimmune diseases, including Covid, multiple sclerosis, Crohn’s disease, and type I diabetes (Supplemental Table 1). Testing for CeD was not performed on these donors, but donors were assumed to be CeD negative for the purpose of comparing CeD-associated TCR prevalence in the context of other autoimmune and infectious diseases.

Of the 1604 total CeD patients, 1339 were adhering to a gluten-free diet at the time of sampling and 265 were consuming gluten normally. We observed no significant difference in sequencing depth (total productive templates) or age between CeD cases and controls. CeD cases skewed significantly more towards female (76%) compared to controls (59%), but this did not result in biased frequency of CeD-specific TCRs (Supplemental Figure 1).

### Published CeD TCRs

We collected a set of 445 published CeD-associated TCRβs from IEDB [30], VDJdb [31] and McPAS [32]. We included only CDR3s with curated V and J genes (rather than imputed). This sequence set included 10 TCRβ sequences binding to epitopes deriving from barley (*Hordeum vulgare*) and the remaining 435 binding to epitopes deriving from common wheat (*Triticum aestivum*).

### Sequencing

Genomic DNA was extracted from frozen, plasma-depleted blood samples using the Qiagen DNeasy Blood Extraction Kit (Qiagen). As much as 18 μg of input DNA was then used to perform immunosequencing of the third complementarity determining (CDR3) regions of TCRβ chains using the ImmunoSEQ Assay (Adaptive Biotechnologies). Briefly, input DNA was amplified in a bias-controlled multiplex PCR, followed by high-throughput sequencing. Sequences were collapsed and filtered to identify and quantitate the absolute abundance (i.e. templates) of each unique TCRβ CDR3 region for further analysis, as previously described [33-35]. To quantify the proportion of T cells out of total nucleated cells input for sequencing, or T cell fraction, a panel of reference genes present in all nucleated cells was amplified simultaneously [36].

### HLA Imputation from TCRβs

HLA genotypes for each subject in the sequence discovery and validation sets (described in the section below) were inferred from the T cell repertoire using TCRβ-based HLA classifiers. Briefly, TCRβ repertoire sequencing from a cohort of ∼40k individuals were assembled. From this large database, TCRβs specific to each of the 145 most common HLAs were identified. These TCRβs were used to build 145 HLA classifiers, one classifier for each HLA [51].

### HLA-aware CeD Enhanced Sequence Discovery

Identification of TCRβs enriched in the CeD patient population was performed using a similar procedure as described previously [34]. Briefly, we first performed HLA inference from TCRβ repertoires as described above, then divided our samples based on presence of one or more of the three main CeD risk alleles (DQ2.5, DQ2.2, and DQ8).

For each risk allele, we assessed occurrence in CeD+ cases and healthy controls for each TCRβ in the sequence discovery set and determined statistical enrichment using a one-sided Fisher’s Exact Test (FET), testing only for enrichment and not depletion in cases. TCRs with an FET p-value below the threshold were defined as Enhanced Sequences (ES). The best FET p-value threshold was determined as 0.002 by evaluating the area under the curve (AUC) of a classifier trained on the clonal breadth (defined below) of ESs in 5-fold cross-validation (Supplemental Figure 2).

We split the TCRβ repertoires into training and validation sets. To reduce genetic biases and ensure generalizability of the classifier across the disease population, the model was trained with genetically matched cases and controls and tested on a validation set coming from two different locations and studies (Table 1). The training set contained 613 CeD-positive cases and 696 CeD-negative controls from the City of Hope study. Of the controls, 588 were first degree and 43 were second degree relatives of CeD subjects in the cohort. The validation set contained 991 CeD-positive cases and 8,436 controls.

**Table 1.**
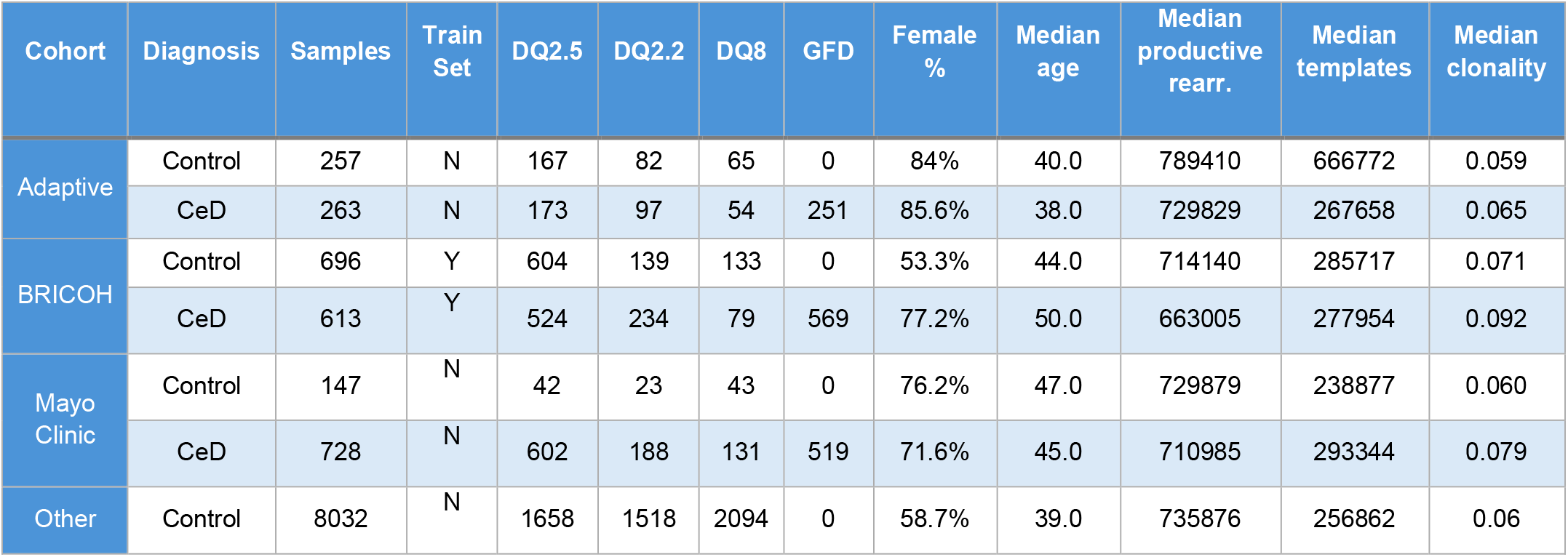
Balance of CeD cases and controls in the training and validation sets.

### Clonal Breadth Analysis

Clonal breadth refers to the number of unique productive TCRβ nucleotide rearrangements corresponding to a given set of sequences in a patient’s repertoire. The unit of the clonal breadth is rearrangements per million (RPM). Since most sequenced repertoires contain fewer than 1 million TCRβs (mean = 210,753), a sampling strategy was employed. Repertoires were randomly down sampled without replacement to 100k TCRβs 10 times to achieve an estimate of RPM. Repertoire with fewer than 100k TCRβs were omitted from analysis.

We used clonal breadth to evaluate the CeD specificity of three sets of TCRβ sequences: our novel set of 318 TCRβs, 409 previously published gliadin-specific TCRβs, and 68 gliadin-specific R-motif TCRβs. Clonal breadth for each of these sequence sets was computed for 884 CeD cases and 3,624 controls from the validation set that harbored at least one of the DQ2 risk alleles (DQ2.5 or DQ2.2).

To assess the significance of ES sets in relation to CeD severity, we carried out correlation analysis between clonal breadth and Marsh score for 367 DQ2+ CeD patients with Marsh score measurements. Marsh score is a metric measuring severity of intestinal damage on the scale of 1, 2, and 3, where 3 is further subcategorized into 3A, 3B, and 3C. A Marsh score of 1 indicates no villous atrophy while 3C indicates total villous atrophy.

### Clonality in CeD Repertoires

To observe the large-scale effects of CeD on the T cell repertoire we computed Productive Simpson Clonality (the square root of Simpson’s diversity index on all productive TCRβ rearrangements), where 1 corresponds to a monoclonal repertoire and 0 a polyclonal repertoire. To account for the consistent increase of clonality with age due to clonal expansion of memory T cells, we partitioned the data into 10-year age bins and performed a Mann-Whitney U (MWU) test within each age bin to assess significance of the difference between cases and controls.

### CeD Associated Viral Activation Analysis

We hypothesized that CeD may stress the immune system and cause a hyper-reactive state, resulting in clonal expansion of TCRs responding to previous viral or other exposures. To evaluate this hypothesis, we divided the subjects based on inferred CMV and EBV positivity, predicted from 26,139 CMV and 9,741 EBV-associated TCRβs from a prior study of over 30k T cell repertoires [36,51]. We then compared clonal breadth associated with CMV and EBV between groups.

### HLA Disease Risk Analysis

HLA enrichment in patients compared to controls was assessed for single and combination alleles using the Fisher’s Exact Test (FET), with Bonferroni correction for multiple comparison. Odd’s ratio (OR) was used to identify additive, non-additive and interaction effects at the HLA loci in CeD patients relative to healthy controls. For a given HLA allele A, let A^+^ denote presence of the allele and A^-^ denote absence. The Odd’s ratio is defined as:

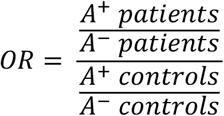

## Results

### CeD HLA Risk Alleles

Before identifying CeD-associated TCRβs, we first inferred HLAs for each subject in the training cohort from their full TCRβ repertoire for the purpose of correcting for HLA bias during in the case and control repertoires (see Methods). We confirmed previously reported HLA effects for different CeD risk alleles [6,38] including relatively equal risk of one copy of the DQ2.5 allele (OR = 6.29, CI = 5.54-7.14) and DQ2.5 in the trans configuration, determined by the DQ7.5/DQ2.2 genotype (OR = 6.70, CI = 5.04-8.88).

Under an additive model, we found strong effects for DQ2.5 (OR = 9.46, 95% CI = 8.22-10.89), DQ8 (OR = 2.40, 95% CI = 1.97-2.92) and DQ2.2 (OR = 2.95, 95% CI = 2.41-3.61) in CeD cases relative to controls. Additionally, we observed previously reported [6] zygosity and interactions effects at these HLA loci (see Table 2). These results demonstrated the feasibility of identifying complex HLA interactions in the HLA loci contributing to Celiac and other autoimmune disease risk [53] using TCRβ repertoires.

**Table 2.** CeD risk conferred by zygosity and interaction effects at the HLA-DQ loci. DQX and DQY indicate a haplotype different from DQ2.5, DQ8, DQ2.2 and DQ7.5.

| Genotype | Reference | OR [95% CI] | p-value |
| --- | --- | --- | --- |
| DQ2.5 | DQX | 9.46 [5.04-8.88] | <0.0001 |
| DQ2.5/DQ2.5 | DQX/DQX | 18.99 [14.09-25.76] | <0.0001 |
| DQ2.5/DQX | DQX/DQY | 6.29 [5.54-7.14] | <0.0001 |
| DQ2.5/DQ2.5 | DQ2.5/DQX | 1.70 [1.44-2.01] | <0.0001 |
| DQ2.5/DQ2.2 | DQ2.5/DQX | 2.08 [1.77-2.42] | <0.0001 |
| DQ2.2 | DQX | 2.95 [2.41-3.61] | <0.0001 |
| DQ2.2/DQ2.2 | DQX/DQX | 1.98 [0.98-3.66] | 0.04 |
| DQ2.2/DQX | DQX/DQY | 2.95 [2.37-3.67] | <0.0001 |
| DQ2.2/DQ2.2 | DQ2.2/DQX | 0.49 [0.26-0.87] | 0.01 |
| DQ8 | DQX | 2.40 [1.97-2.92] | <0.0001 |
| DQ8/DQ8 | DQX/DQX | 5.42 [3.14-9.41] | <0.0001 |
| DQ8/DQX | DQX/DQY | 1.98 [1.60-2.46] | <0.0001 |
| DQ8/DQ8 | DQ8/DQX | 1.09 [0.73-1.62] | 0.60 |
| DQ7.5/DQ7.5 | DQX/DQX | 0.32 [0.10-0.82] | <0.001 |
| DQ7.5/DQX | DQX/DQY | 1.22 [0.97-1.53] | 0.07 |
| DQ7.5/DQ2.2 | DQX/DQX | 6.70 [5.04-8.88] | <0.0001 |

### Enhanced Sequences predict CeD status better than published CeD TCRβs

Using our HLA-aware sequence discovery method (see Methods), we identified a total of 370 CeD enhanced sequences (ESs) from the training cohort (613 CeD cases and 696 controls). The controls included samples with CeD HLA risk haplotypes (DQ2.5, DQ2.2, DQ8 and DQ7.5) to ensure that observed ESs are associated with CeD rather than HLA-linked exposures. The V-gene and J-gene distributions of our ES were similar to published CeD TCRs, with V7-2 being the most common V-gene and J2-7 being the most common J-gene. However, some ES V-genes, such as V2-1, V28-1, V06-05, and V27-1, were not represented in published sequences (Supplemental Figure 7).

Our CeD ESs included 50 TCRβs corresponding to the canonical DQ2.5 CDR3 motif CASSxRxTDTQYF+TRBV7-2/3+TRBJ2-3, known as the R-motif [39] as well as 2 TCRβs previously identified via tetramer sorting, CASSQGSGGNEQFF+TCRBV04-03+TCRBJ02-01 and CASSQGQDTEAFF+TCRBV07-03+TCRBJ01-01, which bind antigens DQ2.5-glia-ω2 and DQ2.5-glia-α1 respectively [40,41]. After removing published sequences, we were left with 318 novel CeD ESs. Comparing the 68 published sequences matching the R-motif and 409 other published TCRβs (see Methods), we observed that the R-motif was significantly more enriched in CeD patients than other published epitope-specific TCRβs (OR 3.45, p=4e-71). However, our novel ESs were far more enriched in CeD than the R-motif TCRs (OR 6.85, p=7e-80) (Table 3), suggesting these novel ESs are better disease biomarkers than what TCR published to date.

**Table 3.**
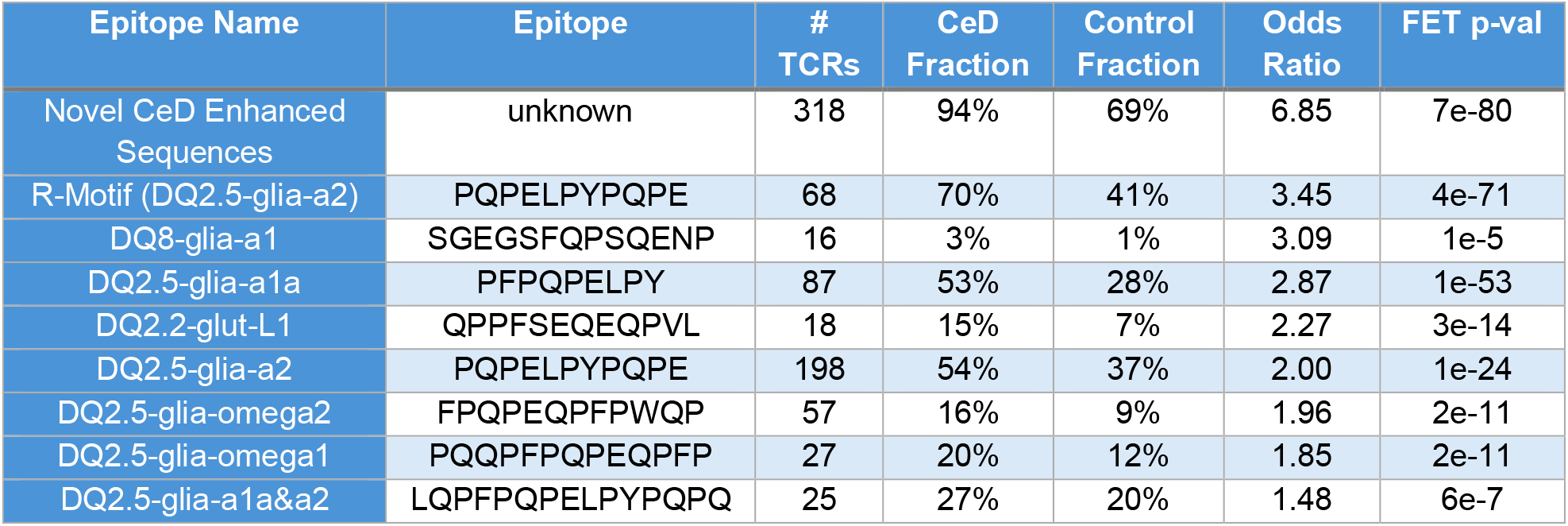
Prevalence of CeD TCRs in validation set grouped by epitope. Odds ratios and corresponding Fisher’s exact p-values indicate enrichment of TCRs in CeD cases versus controls.

We found that our novel ESs could distinguish CeD cases from controls harboring the DQ2 risk allele with significantly higher accuracy (AUC=0.74 GFD, 0.76 non-GFD) than published TCRs (AUC=0.57 GFD, 0.62 non-GFD), including the R-motif (AUC=0.60 GFD, 0.70 non-GFD, Figure 1 top panels). The presence of the risk alleles RPM of novel ESs was significantly elevated in CeD cases compared to controls with viral or bacterial infections (Covid and Lyme) as well as with other autoimmune disorders (type I diabetes, multiple sclerosis, ulcerative colitis and Crohn’s disease) (MWU p<1e-10, Supplemental Figure 3a).

**Figure 1.**
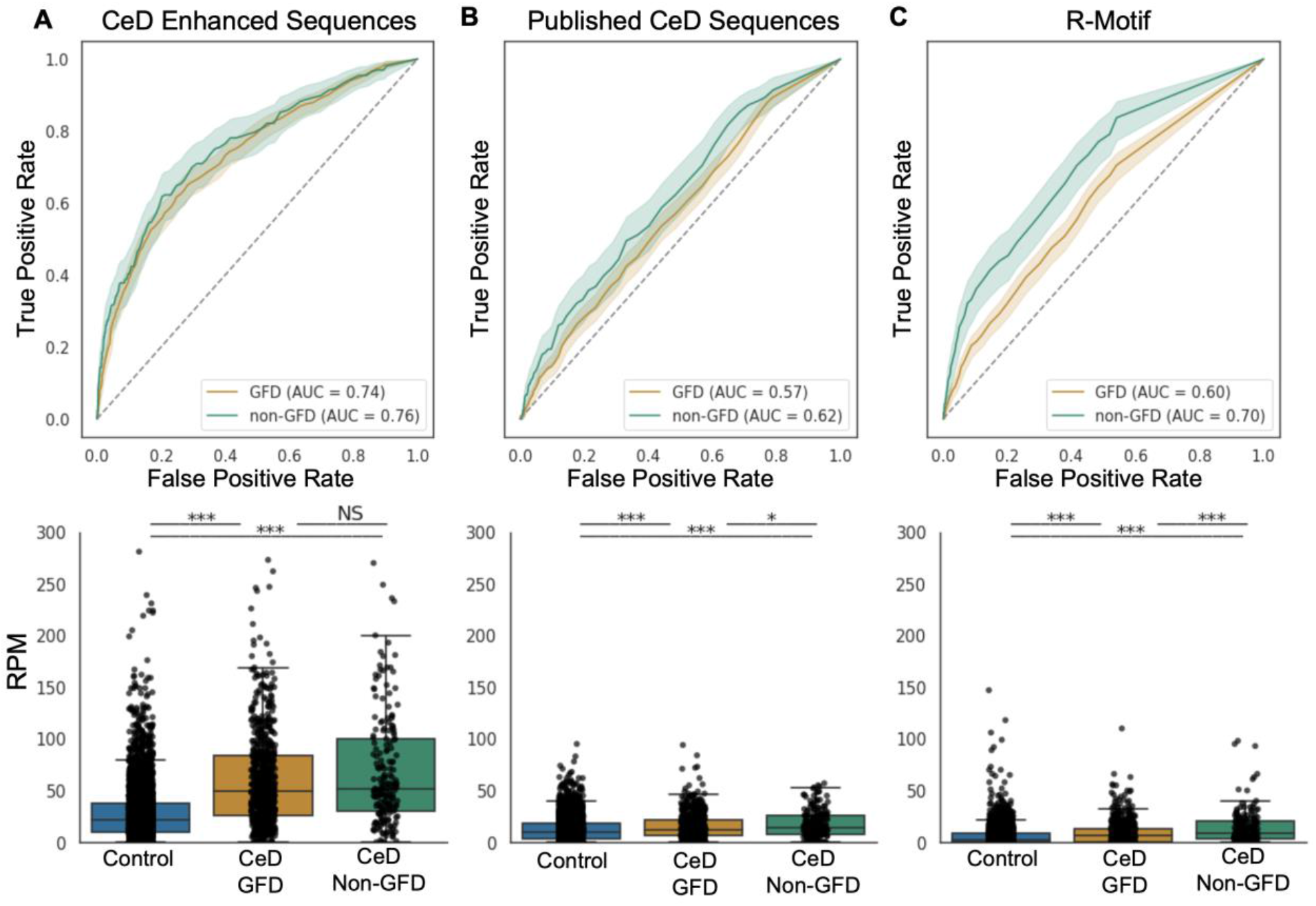
ROC curves (top panels) and clonal breadth (bottom panels) of Enhanced TCRs (N = 318, left panels), published TCRs (N = 409, middle panels), and the R-Motif (N = 68 TCRs, right panels, present in both published and enhanced TCRs) in DQ2+ CeD patients and controls. *p< 0.05, **p< 0.01, ***p< 0.001.

Interestingly, we found no significant difference in the enhanced sequence RPM for patients on or off a gluten-free diet (p >.05), while the RPM of the 409 published CeD-TCRβs and 68 R-motif TCRβs were significantly elevated in patients consuming gluten (R-motif, p<.001, other published, p < 0.05 Figure 1 lower). This suggests the novel CeD ES TCRβs are robust to gluten challenge or absence in the diet.

We note that neither our CeD ESs nor published DQ8-specific TCRs [42] distinguished cases from controls in the DQ8-specific context (Supplementary Figure 4a-b), indicating that the majority of public CeD TCRβs in the peripheral blood bind either DQ2.5 or DQ2.2.

### Enhanced TCR clonal breadth correlated with Marsh score in patients not on GFD

We observed that the clonal breadth of our novel CeD ESs correlated with Marsh score (Pearson r=0.105, p=0.037) (Figure 2a). In comparison, published CeD TCRs (r=0.014, p=0.79) including the R-Motif (r=0.064, p=0.21) were not correlated with Marsh score (Figure 2b-c). This finding suggests that profiling of CeD enhanced sequences in the peripheral blood may be a useful biomarker of severity, disease progression, or recovery.

**Figure 2.**
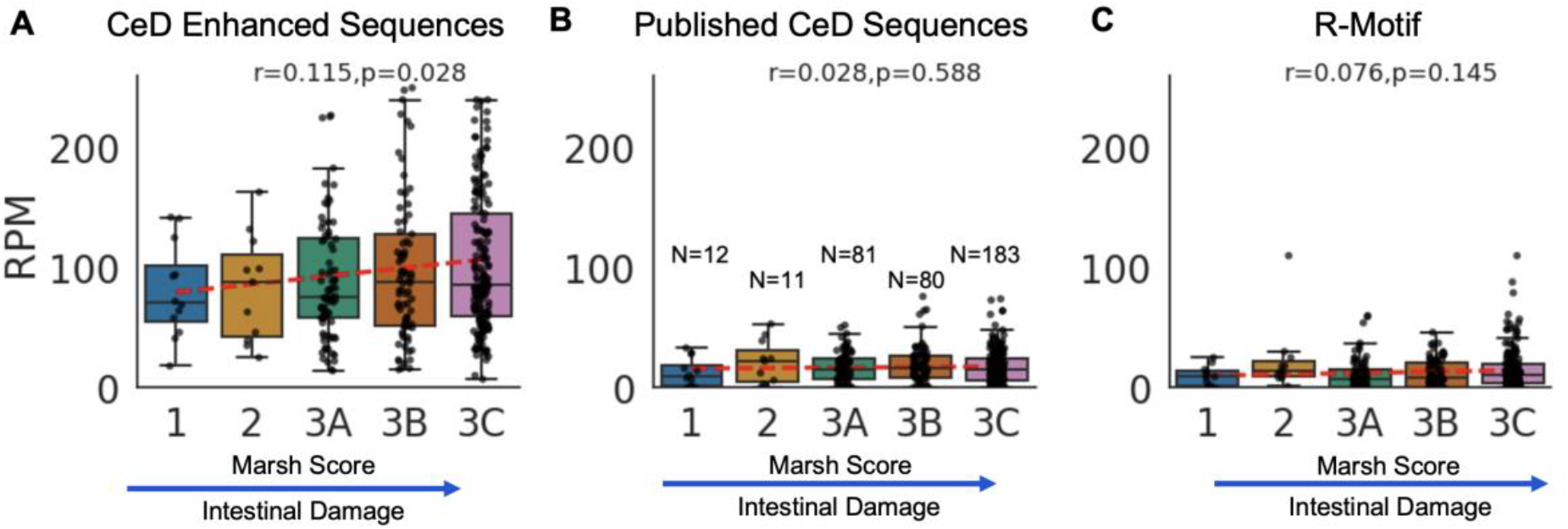
Correlation between Rearrangements Per Million (RPM) and Marsh score. (A) CeD enhanced sequence RPM was positively correlated with Marsh score (Pearson r = 1.15, p = 0.028). (B) CeD published sequence RPM were not significantly correlated with Marsh score. CeD R-Motif sequence RPM was not significantly correlated with Marsh score.

### Higher clonality in CeD repertoires

In addition to measuring the dynamics of CeD-specific TCRβs, we also observed large-scale effects of the disease on T-cell repertoire. We observed a significant increase in Simpson clonality in CeD patients compared to controls. Clonality was significantly increased in CeD cases in all age bins except for bin 70-80 (p<0.001, Figure 3a). This increase in clonality was significant for both CD8+ and CD4+ TCRs (Supplemental Figure 5) and was significant for patients adhering to a GFD (Supplemental Figure 6a) and consuming gluten normally (Supplemental Figure 6b).

**Figure 3.**
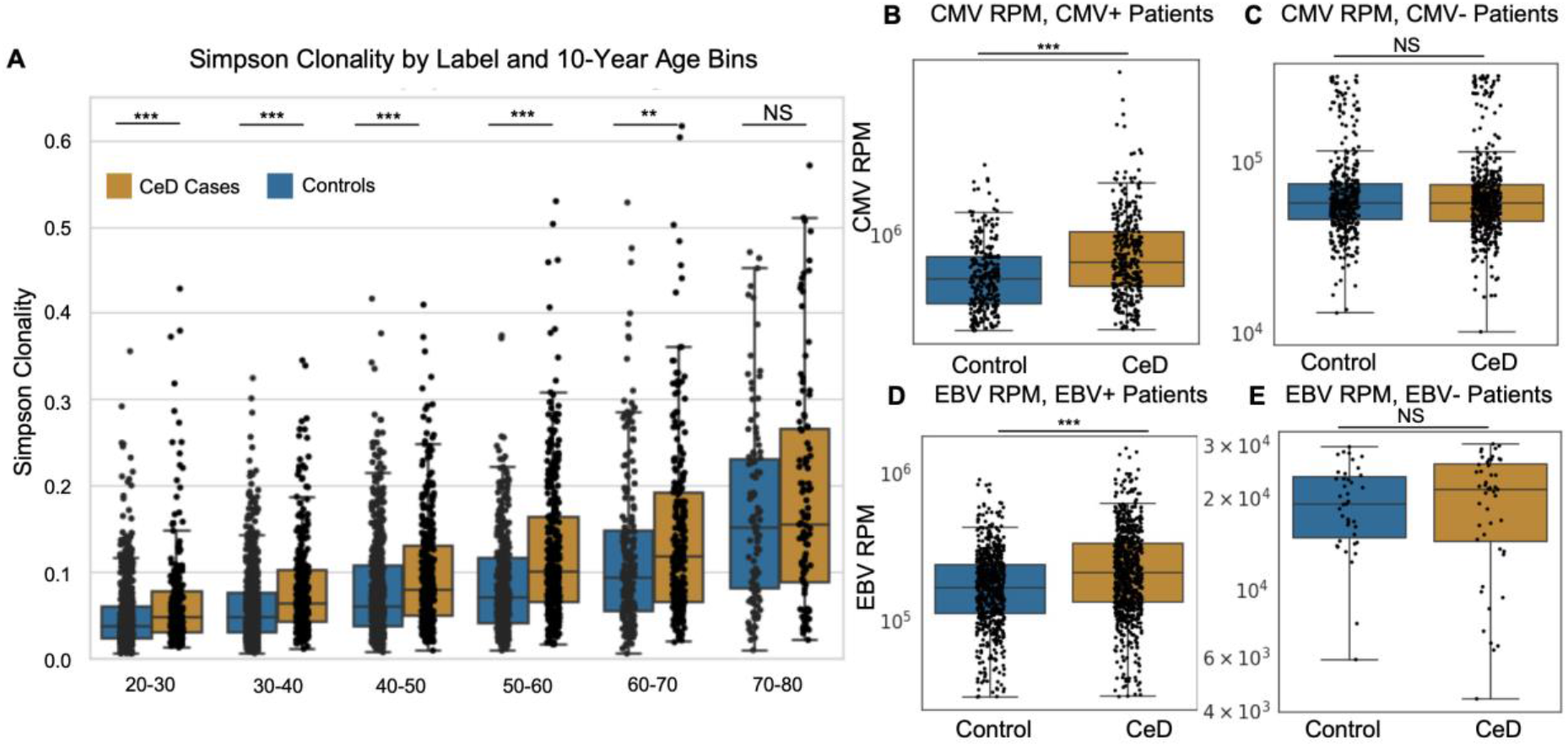
(A) Simpson clonality between CeD cases (orange) and controls (blue) grouped in 10-year age bins. (B) CMV RPM between CMV-positive CeD cases and controls. CeD cases had significantly elevated CMV RPM versus controls (MWU p<1e-9) (C) CMV RPM between CMV-negative CeD cases and controls. CeD cases did not significantly differ from controls (p>.6). (D) EBV RPM between EBV-positive CeD cases and controls. CeD cases had elevated EBV RPM versus controls (MWU p<1e-13). (E) EBV RPM between EBV-negative CeD cases and controls. CeD cases did not significantly differ from controls (MWU p>.29). *p<.05, **p<.01, ***p<.001.

### Expansion of T cells from viral exposures

We confirmed that CeD patients positive for CMV and EBV had significantly higher CMV and EBV RPM respectively than CMV and EBV-positive controls (p < 1e-9, p<1e-13 respectively, Figure 3b,d). This indicated an expansion of viral memory T cells in subjects with CeD. As expected, we observed no difference in CMV and EBV RPM for CMV- and EBV-patients. Importantly, we note that the inferred rate of positivity for these viral infections was not increased in CeD cases compared to controls, with 43% of CeD cases and 44% of controls being CMV-positive and 94% of CeD cases and 94% of controls being EBV positive (p >.5). This result indicated that CMV and EBV infection are not likely to be necessary triggers for CeD, which differs from other autoimmune disorders such as multiple sclerosis, where EBV infection was a prerequisite for disease onset [45].

## Discussion

Through immune sequencing of the TCRβ repertoires from the peripheral blood, we identified over 300 novel public TCRβ enhanced sequences (ESs) which were present in significantly higher proportion in 696 CeD cases versus 613 controls. We demonstrated the utility of these TCRβs in a validation set consisting of 884 DQ2+ CeD cases and 3,624 DQ2+ controls. The novel sequence set significantly outperformed published TCRβs in distinguishing CeD cases from controls, especially for subjects adhering to a gluten-free diet. Most novel TCRβs had similar V-gene usage patterns as published ones (V7-2, V20-1), while some novel TCRβs had previously unobserved V-gene usage patterns (V27-1, V06-5, V28-1, V2-1). Additionally, the ESs ability to distinguish cases from controls despite sharing the CeD risk haplotypes further confirms that these sequences are primarily CeD driven and not associated with a specific HLA haplotype.

The discovery of novel public ESs that are robust to dietary gluten intake suggests that these TCRs may recognize antigens that are central to the pathogenesis of CeD but have remained elusive in previous studies. One possibility is that these ESs recognize non-gluten antigens. Another possibility is that they recognize gluten antigens but harbor a phenotype that promotes longer persistence in memory than other gluten-reactive T cells, such as those with the R-motif. Additionally, the identification of these TCRs in the peripheral blood could indicate a systemic immune response that transcends the localized gut inflammation typically associated with CeD. This finding opens new possibilities for exploring the role of systemic autoimmunity in CeD that may have implications for the identification of novel therapeutic targets that can modulate this immune response. However, this study was limited to only assessing the statistical association of TCRβs to CeD. Future studies will be needed to experimentally confirm the antigen-specificity of these public ESs to truly understand their relationship to disease pathology.

We further observed that the clonal breadth of these novel ESs correlated with increasing intestinal damage as measured by Marsh score. These public TCRβs could potentially serve as a non-invasive surrogate marker for assessing intestinal damage, potentially reducing the need for repeated biopsies. However, follow-up studies would be required to validate and calibrate this signal. The lack of correlation observed with other published sequences, including the gliadin-a2-binding “R-motif”, underscores the unique nature of these novel ESs and suggests that they may be capturing aspects of the disease process that are not reflected in previously identified TCR sequences. These findings demonstrate that public TCRβs in the peripheral blood are important biomarkers for CeD and highlight the advantage of unbiased sequence discovery through non-invasive bulk immune repertoire sequencing of case/control cohorts. This experimental design enables robust detection of autoimmune-associated TCRs without restricting to specific antigens.

We additionally presented evidence of increased repertoire clonality and clonal expansion of viral TCRβs in CeD. We believe this could have important implications for understanding the mechanisms underpinning autoimmunity in CeD. One mechanism that could explain the expansion of memory CMV and EBV TCRs and overall higher clonality is bystander activation. During bystander activation, self-reactive T cells induce activation of other antigen-specific memory T cells to produce inflammatory cytokines [43-45]. Another possible mechanism is viral reactivation. This occurs when a latent infection switches to a lytic stage of replication, resulting in an adaptive immune response [46]. CMV and EBV both have latent stages, where they remain dormant in infected cells. T cell reactivation associated with viruses has been studied in the context of autoimmunity [52], though prior work has not implicated CMV or EBV reactivation in CeD [47-49].

The hypothesis that bystander activation or viral reactivation could be contributing to the observed increase in clonality aligns with emerging evidence in other autoimmune diseases, where viral infections have been implicated as triggers for disease onset or flares. More research is needed to understand the role of latent viral infections and autoimmunity, especially in CeD. Future studies could focus on longitudinal monitoring of CeD patients with known viral exposures to assess changes in TCR repertoire and symptom severity over time.

While a gluten-free diet remains the standard of care for CeD management, many therapeutics are in active development. Our novel CeD public TCRβs possess the following desirable qualities 1) measurable with non-invasive technology, 2) robust to diet and likely associated to a broad set of celiac-associated antigens 3) strongly correlated with disease severity that can only be measured with invasive endoscopy. Furthermore, our observation of increased CMV and EBV clonal breadth in CeD patients highlights the usefulness of bulk TCR sequencing for monitoring viral reactivation in CeD, which may prove to be important for symptom management.

Together, these properties suggest bulk TCR sequencing, and particularly public CeD TCRs, may be ideally suited in assessing treatment outcome in clinical trials.

## Author Contributions

JAM, SLN, RSC, EVM, GB, and PG designed the initial study. RE and RB performed data analysis with support from WZ and HCH. BH and HSR oversaw generation of the T-cell receptor sequencing data. RE, RB, JAM, RSC, and EBM wrote the manuscript, with contributions from SLN, WZ, and HC. All authors commented on and approved the final version of the manuscript.

## Author Declarations

RE, RB, WZ, HCH, BH, and HSR have employment and equity ownership with Adaptive Biotechnologies.

## Data Availability

All clonal breadth and model scores used to generate figures will be made available as a supplementary file. Full TCR repertoire sequencing data for 520 individuals sequenced in this study will be made available at https://clients.adaptivebiotech.com/pub/elyanow-2024-s.

