## Supplementary Figures and Tables for "Shared TCRs in peripheral blood offer robust celiac disease classification independent of gluten Intake"

### Supplemental Figures and Tables

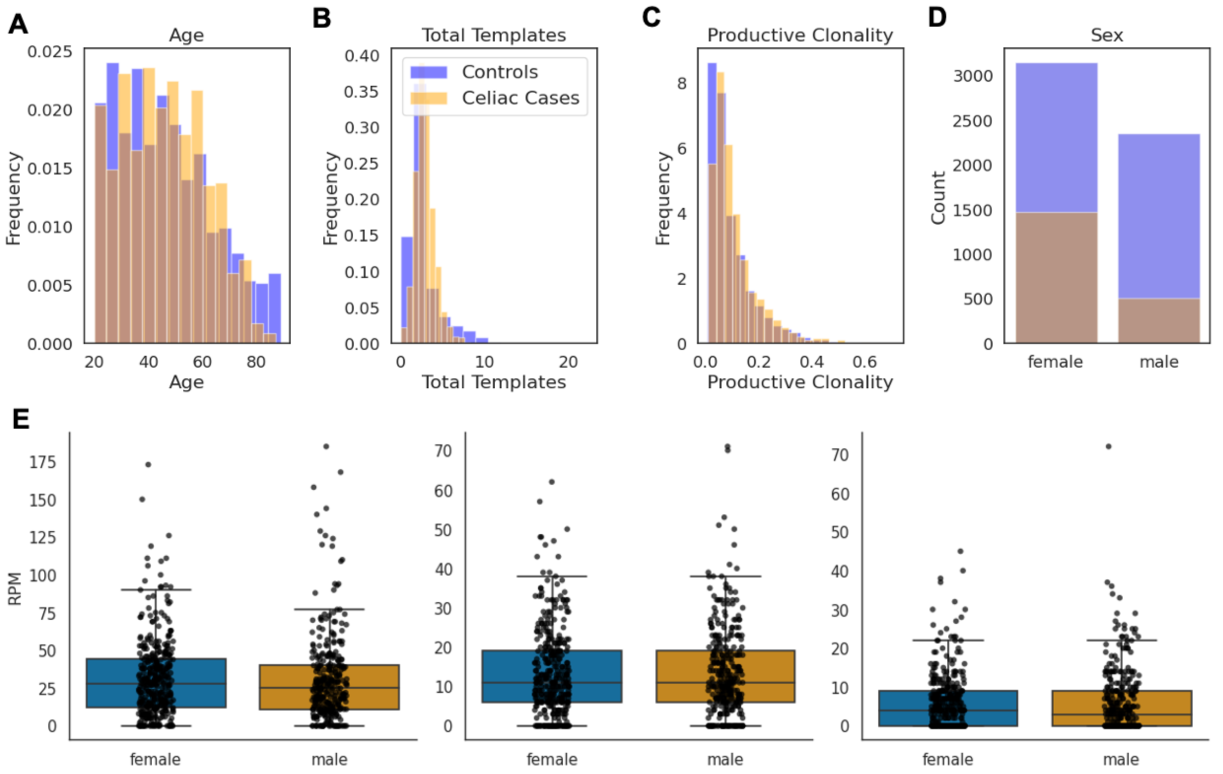


Supplementary Figure 1: Basic QC/metadata. (A) Age distribution of cases (orange) and controls (blue). (B) Total productive templates distribution. (C) Productive clonality distribution. (D) Sex distribution. (E) Repertoires Per Million (RPM) of CeD enhanced TCRs (left), published CeD TCRs (middle) and R-motif (right) does not differ between males and females.

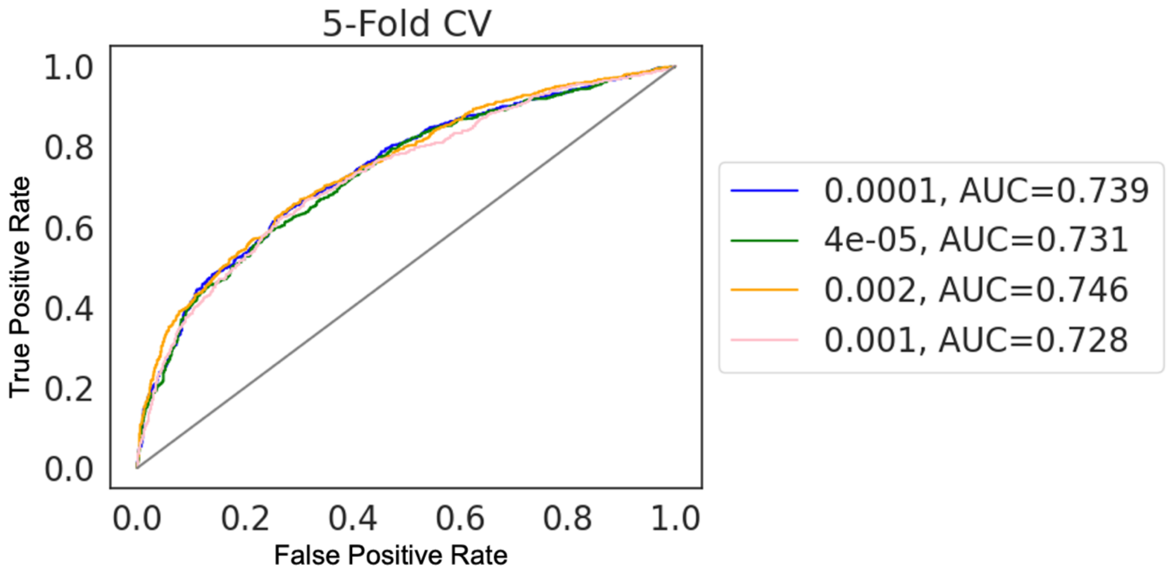


Supplementary Figure 2: 5-fold CV in training to decide p-value cutoff for Fishers-Exact Test. All p-values within range performed well, p=0.002 was selected.

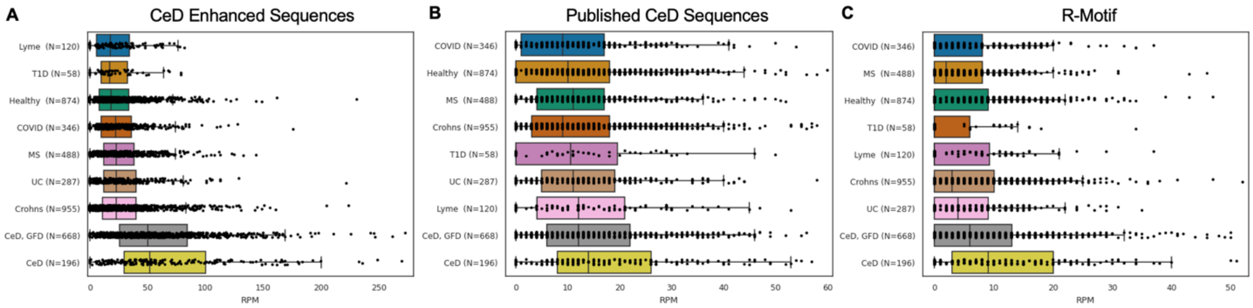


Supplemental Figure 3. RPM broken down by disease for (A) CeD enhanced sequences (B) published CeD sequences, and (C) R-Motif sequences.

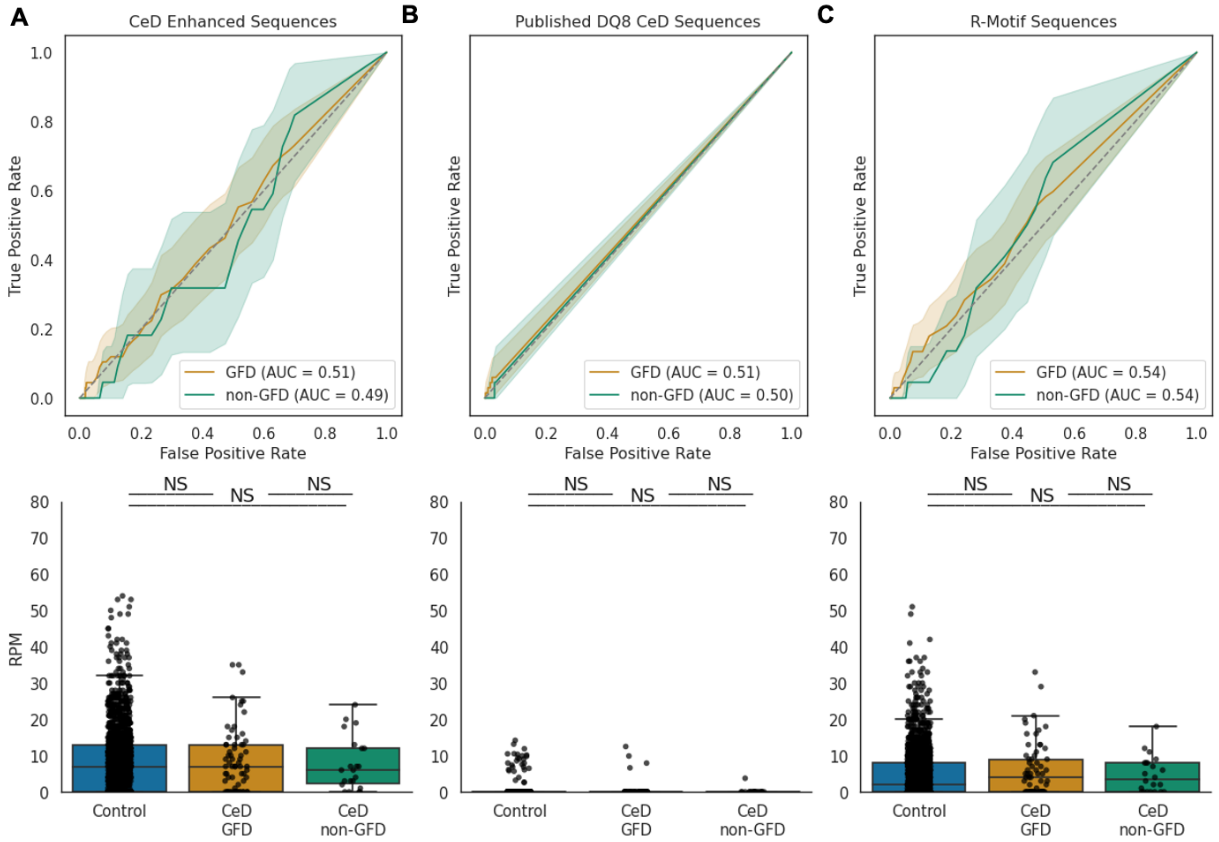


Supplemental Figure 4. RPM of Enhanced TCRs, published DQ8 TCRs, and the R-Motif in DQ8+/DQ2- Celiac patients and controls. No significant enrichment of CeD sequences in cases vs. controls in GFD or non-GFD context for any sequence set ($p > .05$).

| Disease | Samples | DQ2.5 | DQ2.2 | DQ8 | Female % | Median age | Median templates | Median clonality |
|---|---|---|---|---|---|---|---|---|
| **Healthy** | 2698 | 1080 | 495 | 682 | 62% | 39 | 200285 | .054 |
| **Covid** | 823 | 169 | 181 | 213 | 44% | 60 | 352003 | .148 |
| **Lyme** | 301 | 58 | 49 | 87 | 43% | 53 | 203524 | .062 |
| **T1D** | 106 | 48 | 11 | 49 | 62% | 28 | 175376 | .029 |
| **Crohn's** | 2213 | 398 | 488 | 562 | 57% | 37 | 332322 | .067 |
| **UC** | 844 | 137 | 126 | 184 | 51% | 40 | 294348 | .056 |
| **MS** | 1224 | 263 | 204 | 301 | 76% | 37 | 405180 | .057 |

Supplemental Table 1. Breakdown of validation control samples by disease.

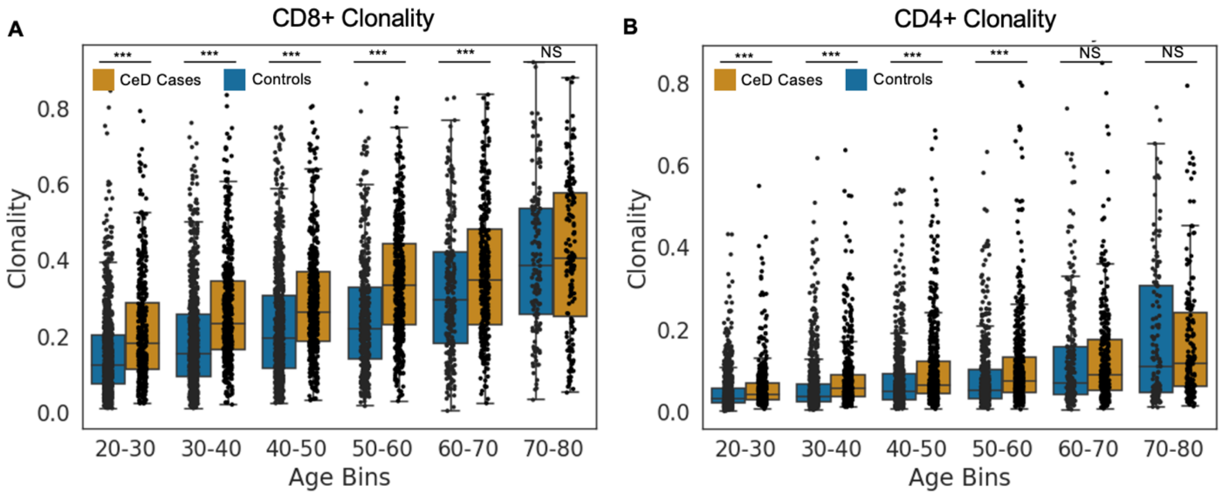


Supplemental Figure 5. (a) CD8+ clonality between CeD cases (orange) and controls (blue) grouped in 10-year age bins. (b) CD4+ clonality between CeD cases and controls. *p< .05, **p< .01, ***p< .001.

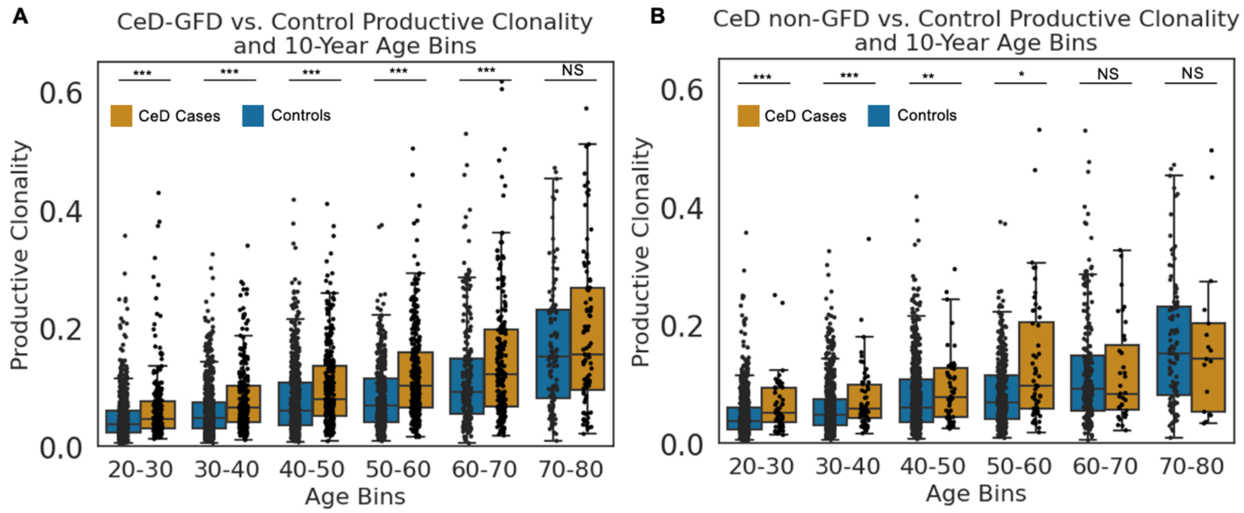


Supplemental Figure 6. (A) Productive clonality between CeD patients on a gluten free diet (orange) and controls (blue) grouped in 10-year age bins. (B) Productive clonality between CeD patients on a regular diet and controls, grouped in 10-years age bins *p< .05, **p< .01, ***p< .001.

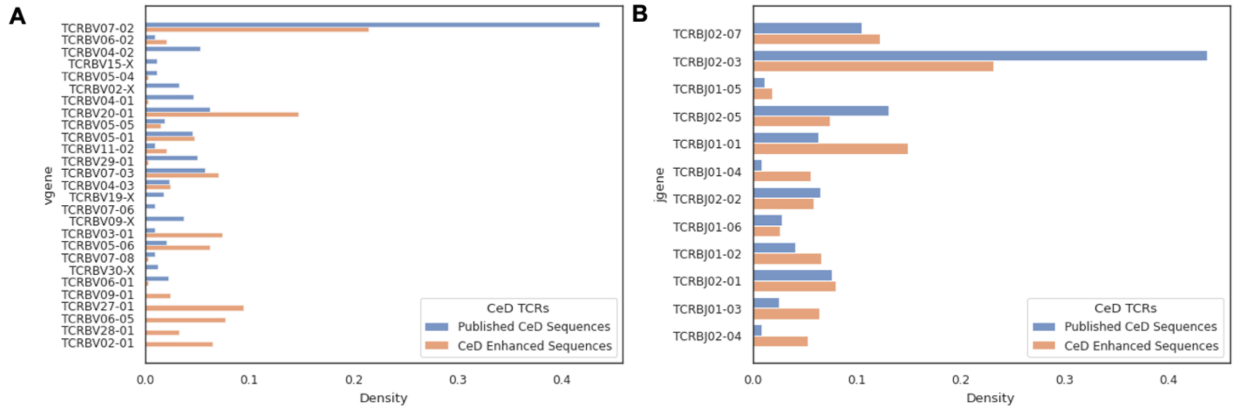


Supplemental Figure 7 (A) V-gene and (B) J-gene density for published CeD Sequences (blue) and novel CeD enhanced sequences (orange).
